# Virus-against-virus dominant-negative interference strategy targeting a viral CC chemokine prevents cytomegalovirus-related neurodevelopmental pathogenesis

**DOI:** 10.1101/2025.02.11.637423

**Authors:** Sylvian Bauer, Sarah Tarhini, Emmanuelle Buhler, Saswati Saha, Thomas Stamminger, Daniel N. Streblow, Nail Burnashev, Hervé Luche, Pierre Szepetowski

## Abstract

**Background:** Congenital cytomegalovirus (CMV) infections are one leading cause of human neurodevelopmental disorders. Increasing evidence for the pathogenic involvement of brain immune alterations was obtained in the recent years. Host and virus-encoded chemokines might play important roles in CMV-related neuropathogenesis by regulating leukocyte trafficking and microglia recruitment in the CMV-infected brains, and by interfering with key neurodevelopmental steps. In a rat model of CMV infection of the fetal brain *in utero* that leads to detrimental neurologic and other severe phenotypes postnatally, we reported on the early alteration of microglia and on the infiltration of the infected brains by lymphoid and myeloid cells. Particularly, expression of the *r129* gene encoding the viral chemokine RCK3 was detected as early as 24h post-infection; together with the previously reported chemotaxis properties exerted by RCK3 on lymphocytes and on macrophages *in vitro*, this suggested that RCK3 might be involved in the brain immune cell alterations seen in the CMV-infected developing brains, and in the related neuropathogenesis.

**Methods:** Infection of the rat fetal brain was done by intracerebroventricular injections *in utero* of either rat CMV encoding wild-type (wt) RCK3 (RCMV-wt), or a mutant CMV counterpart encoding RCK3 with a deletion in its chemokine domain (RCMV-r129ΔNT). As RCMV-r129ΔNT had shown dominant-negative effects on the chemotaxis properties of RCK3 *in vitro*, simultaneous and successive co-infection rescue assays were also performed. The detrimental postnatal phenotypes *in vivo* and the epileptiform activity *ex vivo* usually detected after infection of the rat developing brain with RCMV-wt were monitored in the RCMV-r129ΔNT condition and in co-infection assays.

**Results:** In sharp contrast with RCMV-wt whose infection of the fetal brain led to decreased postnatal survival, impaired sensorimotor development, hindlimb hyperextension and epileptic seizures in neonatal pups, RCMV-r129ΔNT infection was not associated with any severe postnatal phenotype *in vivo*. Consistently, the epileptiform activity recorded in most neocortical slices from RCMV-wt-infected pups was not detected in any slice from RCMV-r129ΔNT-infected pups. Simultaneous co-infection assays led to dramatic prevention against the postnatal phenotypes *in vivo* and the altered network activity *ex vivo*, revealing a dose-dependent rescuing effect exerted by RCMV-r129ΔNT on RCMV-wt. Importantly, successful rescue was also obtained when the mutant RCMV-r129ΔNT was inoculated in the fetal brains either before or after infection with RCMV-wt.

**Significance:** Our data demonstrate the crucial neuropathogenic role of RCK3 CMV chemokine *in vivo*. The dramatic success and apparent safety of the dominant-negative RCK3 rescue assays *in vivo* provide a proof-of-principle for the beneficial use of a virus-against-virus approach against CMV-related pathogenesis.

## Introduction

Besides genetic defects, a large proportion of neurodevelopmental disorders is caused or at least influenced by exposure to various environmental insults during pregnancy. These include infections by pathogens such as rubella virus, zika virus, or cytomegalovirus (CMV) - a member of the *Herpesviridae* family. Generally, congenital infections cause morbidity and mortality throughout the world and are of considerable public health impact. As a matter of fact, congenital CMV infections (0.2 to 2% of all live births worldwide, 0.5% in Western Europe) represent one leading cause of human neurodevelopmental disorders. 15-20% of neonates infected with CMV during pregnancy are born with, or will develop, severe neurologic and other defects (e.g. growth retardation, brain malformations, hearing impairment, cerebral palsy, epileptic seizures, intellectual disability) (Adler & Nigro, 2013; Leruez-Ville et al., 2020; Manicklal et al., 2013). Furthermore, an influence on the risk to neuropsychiatric diseases such as schizophrenia or autism spectrum disorders has been proposed (Børglum et al., 2014; Leruez-Ville et al., 2020; Turriziani Colonna et al., 2020).

Despite the incidence and the worldwide medical and socioeconomical burden of congenital CMV, the pathophysiological mechanisms underlying the emergence of neurodevelopmental disorders have long remained elusive. No satisfactory preventive or curative therapy is available so far. Whereas the pathophysiology of congenital CMV disease is inherently complicated and involves several maternal and fetal steps, insights into the early events following CMV infection of the developing brain are particularly needed. CMVs are species-specific; thus, relevant animal models are critical to the understanding of the mechanisms involved in CMV congenital brain disease (Cekinovic et al., 2014; Schleiss, 2006). Convergent insights into the alteration of innate and adaptive immune responses in the infected brains have emerged from such models (Bradford et al., 2015; Cloarec et al., 2016; Mutnal et al., 2011; Sakao-Suzuki et al., 2014; Seleme et al., 2017; Slavuljica et al., 2015; Zhou et al., 2022). Recent findings in rats and mice have emphasized the crucial neuropathophysiological influence exerted in CMV-infected developing brains by inflammatory processes (Kosmac et al., 2013; Tarhini et al., 2024) and by the recruitment of various immune cells such as memory CD8^+^ T-cells (Brizić et al., 2018), natural killer and type 1 innate lymphoid cells (Kveštak et al., 2021), as well as by the alteration of fetal microglia (the resident immune cells of the brain) (Cloarec et al., 2018). Analyses of CMV-infected human fetal brains also showed a significant association between the severity of brain alterations on the one hand, and the higher detection of innate immune cells on the other hand (Sellier et al., 2020).

Owing to their role in cell migration and recruitment, chemokines, a subgroup of small cytokines with chemoattractant properties, might influence the infiltration of the developing brain by monocytes-derived macrophages and by lymphocytes as well as the modification of microglia phenotype that can be seen in the context of CMV infection. Based on the number and spacing of conserved N-terminal cysteines, chemokines can be classified into one of four different categories (namely C, CC, CXC, or CX3C) (Hughes & Nibbs, 2018). CMV interferes with the host chemokine system (Luganini et al., 2016; Cheeran et al., 2001); this might lead to central nervous system dysfunction and impaired behavior later in life, as chemokines influence critical stages of fetal brain development (Deverman & Patterson, 2009). Furthermore, the large (about 235 kilobases in size), double-stranded CMV genome encodes viral chemokines and chemokine receptors (Pontejo & Murphy, 2017) that may also participate in CMV-related neuropathogenesis during development. Amongst the two or more chemokines expressed by most CMVs (Pontejo & Murphy, 2017), human CMV UL128, mouse CMV MCK-2 and rat CMV RCK3 are CC chemokines encoded in a syntenic region of their respective genomes and are considered functionally homologous, despite sharing low amino acid sequence identities (Pontejo & Murphy, 2017; Vomaske et al., 2012). In addition to their chemotaxis properties, both UL128 and RCK3 CMV chemokines may act as direct actors of viral cell entry as parts of the respective human and rat CMV pentameric complexes (Chandramouli et al., 2017; Jones et al., 2020). Particularly, RCK3 is a secreted protein encoded by the *r129* gene in the rat CMV genome. It promotes the migration of rat macrophages and T cells *in vitro* (Vomaske et al., 2012). Cell attraction driven by RCK3 is dependent on its N-terminal CC chemokine domain, as various mutations and short deletions targeting this amino acid motif led to chemotaxis impairment while not decreasing virus growth and production in infected cells (Vomaske et al., 2012). In contrast, defects in the C-terminal domain of RCK3 impair cell entry, but not chemotaxis properties (Jones et al., 2020; Vomaske et al., 2012). Two RCK3 isoforms bearing mutations of the CC chemokine motif were shown to exert a dominant-negative effect on the chemotaxis properties of wild-type (wt) RCK3 (Vomaske et al., 2012).

We previously reported on a model of CMV infection of the rat fetal brain *in utero*, in which the infiltration by monocytes and lymphocytes and the alteration of microglia in the fetal and neonatal brains were associated with a series of postnatal manifestations that recapitulated key neurologic and other features of human congenital CMV (Cloarec et al., 2018; Tarhini et al., 2024). At the molecular level, expression of *r129* was detected in the fetal brain as early as 24 h after CMV inoculation at embryonic day 15.5 (E15.5), and then increased with time (Cloarec et al., 2016). Such an expression pattern, and the chemotaxis properties of the *r129*-encoded RCK3 protein, raised the possibility that RCK3 participated in the early stages of CMV-related pathogenesis in the developing brain. Here, using a rat CMV strain (RCMV-wt) encoding wt RCK3 and a genetically-engineered derivative strain (RCMV-r129ΔNT) encoding a mutant, dominant-negative RCK3 isoform with a short, 10 amino acid deletion in the CC domain (RCK3-ΔNT) (Vomaske et al., 2012), we have addressed the possible role of RCK3 in CMV-related developmental neuropathogenesis *in vivo*.

## Material and Methods

### Ethical statement

Animal experimentations were performed in accordance with the French legislation and in compliance with the European Communities Council Directives (2010/63/UE). This study was approved under the French department of agriculture and the local veterinary authorities by the Animal Experimentation Ethics Committee No. 014 under licenses No. APAFIS#7256-2016100715494790 v3 and APAFIS#21469-2019071211017635 v2.

### CMV infections in utero

Naive timed pregnant Wistar rats were obtained at Janvier Labs (France) and kept at room temperature under conditions of optimal health and hygiene at INMED animal facility. Two recombinant rat CMV Maastricht strains were used: RCMV-wt encodes wt RCK3 chemokine and harbors a green fluorescent protein (GFP) expression cassette; mutant RCMV-r129ΔNT was derived from the RCMV-wt strain mentioned above and encodes a mutant RCK3 isoform (RCK3-ΔNT) with a 10-aminoacid deletion in its CC chemokine motif (Vomaske et al., 2012). Strains productions, purifications and titrations were done as reported previously (Vomaske et al., 2012). *In utero* icv injections were performed in embryos from timed pregnant rats that were anaesthetized with isoflurane (4% for induction, then 2.5%) at embryonic days 14.5 (E14.5), 15.5 (E15.5) or 16.5 (E16.5), depending on the experimental procedure and as previously reported (Cloarec et al., 2016, 2018; Tarhini et al., 2024). In each embryo, 1 μl of minimal essential medium (MEM) with FastGreen (2 mg/ml; Sigma-Aldrich, Saint-Quentin Fallavier, France) and 5% fetal calf serum, either containing 3.5×10^3^ plaque forming unit (pfu) of rat CMV of interest (RCMV-wt or RCMV-r129ΔNT), or not (control), was injected icv at the appropriate embryonic day *via* pulled glass capillaries and a microinjector (PV 820 Pneumatic PicoPump, World Precision Instruments, Friedberg, Germany). For the *in vivo* phenotypic evaluations of simultaneous co-infection experiments with RCMV-wt and RCMV-r129ΔNT at E15.5, procedure was identical except that 1.75×10^3^ pfu of each RCMV was injected.

### Phenotyping of pups

Postnatal phenotyping was done on a daily basis in the first postnatal weeks, based on our previous observations on CMV-infected pups (Tarhini et al., 2024). In addition to the overall assessment of the pups and to the monitoring of body weight, success to the classical developmental righting and cliff aversion sensorimotor tests, and the presence of hindlimb hyperextension and of generalized tonic-clonic epileptic seizures (GTCS), were monitored as previously done (Cloarec et al., 2018; Tarhini et al., 2024). Hindlimb hyperextension was detected visually in pups that had a postural misplacement and immobility of their hindlimbs. GTCS were detected visually, usually after animal handling, especially during cage changing and behavioral testing. GTCS consisted in a classical behavior sequence including movement arrest and loss of postural equilibrium, then hypertonic posture of the trunk, limbs and tail, symmetrically, and then repeated, large clonic movements of all limbs, often with respiratory arrest, incontinence, motor automatisms such as chewing and grooming, terminated by a catatonic phase.

### Extracellular electrophysiological recordings

Recordings were performed following the same experimental procedure as previously reported (Tarhini et al., 2024). Briefly, for slices preparation, brains were rapidly removed from P14-P16 rat pups euthanized after isoflurane (3-4%) anesthesia and placed in an ice-cold slicing solution containing (in mM): 126 choline chloride, 2.5KCl, 1.25 NaH_2_PO_4_, 7 MgCl_2_, 0.5 CaCl_2_, 26 NaHCO_3_, 10 D-glucose, bubbled with 95% O_2_/5% CO_2_. 300 μm-thick coronal neocortical slices (Vibratome Leica VT1000S, Leica Microsystems Inc., USA) were left to recover in artificial cerebrospinal fluid solution (ACSF) containing (in mM): 125 NaCl, 3.5 KCl, 1 CaCl_2_, 2 MgCl_2_, 1.25 NaH_2_PO_4_, 26 NaHCO_3_ and 10 glucose, equilibrated at pH 7.3 with 95% O_2_ and 5% CO_2_ at 33°C for about 30 min, then at room temperature (22-25°C) for about an hour. The final osmolarity of the ACSF solution was 300 ± 5 mOsm. Slices were then transferred to the recording chamber and perfused with oxygenated ACSF recording solution at 3 mL.min^-1^ at 35°C under a Leica DMLFS microscope (Leica Microsystems Inc., USA). Electrophysiological extracellular recordings of spontaneous local field potentials were obtained from layer IV of the neocortex using a coated nichrome electrode (California Fine Wire, Grover Beach, CA, USA) with a low-noise multichannel DAM-80A amplifier (WPI, UK; high-pass filter 3Hz; low-pass filter 1 kHz; gain x1000). Data was digitized using a Digidata 1322A and recorded using Axoscope (Molecular Devices, CA, USA). Recordings were filtered with a low pass 500 Hz filter for analysis using Clampfit software (Molecular Devices, USA). Spikes’ frequency was estimated using the threshold event detection with threshold set as twice the background noise.

### Statistics

Data were expressed as means ± s.e.m. (standard error of the mean). Non-parametric Kruskall-Wallis test followed by Dunn’s post-hoc test for multiple comparisons (data sets > 2) was used wherever appropriate. Fisher’s exact test, with pairwise comparisons and Benjamini-Hochberg correction if needed, was used to assess the overall risks to hindlimb hyperextension and to GTCS, to compare the number of pups exhibiting ictal-like events in extracellular slice recordings, and to verify the lack of sex ratio differences between cohorts. The statistical analyses and representation were performed using R version 4.3.1 (R Core Team: The R Project for Statistical Computing, R Foundation for Statistical Computing, Vienna, Austria; https://www.r-project.org/), Rstudio version 2023.06.2+561 ‘Mountain Hydrangea’ release (RStudio: Integrated Development Environment for R. Posit Software, PBC, Boston, MA, USA; http://www.posit.co/) and GraphPad Prism version 10.1.1 (GraphPad Software, San Diego, California, USA). Survival analysis was performed using the Kaplan-Meier method, with the survival curves among treatment groups compared using the Log-Rank test. The “survival” and “survminer” packages in R were utilized for survival analysis (Therneau & Grambsch, 2000). The “lsmeans” package was employed for deriving least-squares means and contrasts (Lenth, 2016), while the “lme4” package was used for the fitting of mixed models for body weight gain and for the righting reflex and cliff aversion (Bates et al., 2015). Scripted R codes used for the analyses are available upon reasonable request. Significance thresholds were set at p < 0.05.

## Results

### Mutant RCMV encoding RCK3 with a deleted CC chemokine motif is not pathogenic

Rat CMVs with a GFP expression cassette (Vomaske et al., 2012) were injected intracerebroventricularly (icv) in embryos from timed pregnant rats at embryonic day 15.5 (E15.5), as previously done (Figure 1A) (Cloarec et al., 2016, 2018; Tarhini et al., 2024). To start in evaluating the possible role of RCK3 in CMV-related neurodevelopmental pathogenesis, mutant CMV encoding RCK3 with a 10-amino acid deletion in its chemokine domain (RCMV-r129ΔNT) (Figure 1B) was injected into the cerebral ventricles of rat embryos, using the exact same experimental protocol as with RCMV-wt. Consistent with our previous reports (Cloarec et al., 2018; Tarhini et al., 2024), decreased survival and body weight gain, failure to perform the righting reflex and cliff aversion tests taken as classical readouts of proper sensorimotor development, and the occurrence of hindlimb hyperextension and of generalized tonic-clonic seizures (GTCS), were all seen in the three first postnatal weeks in cohorts of pups submitted to icv injections of RCMV-wt (Figure 1C-H). Remarkably, pups infected *in utero* with RCMV-r129ΔNT did not show any of the deleterious phenotypes seen with RCMV-wt, apart from slight differences in body weight gain and in the success to cliff aversion test in the very first postnatal days (Figure 1C-H). The lack of any severe phenotype *in vivo* upon infection with RCMV-r129ΔNT suggested that viral chemokine RCK3 may play an important role in CMV-related neuropathogenesis.

**Figure 1.**
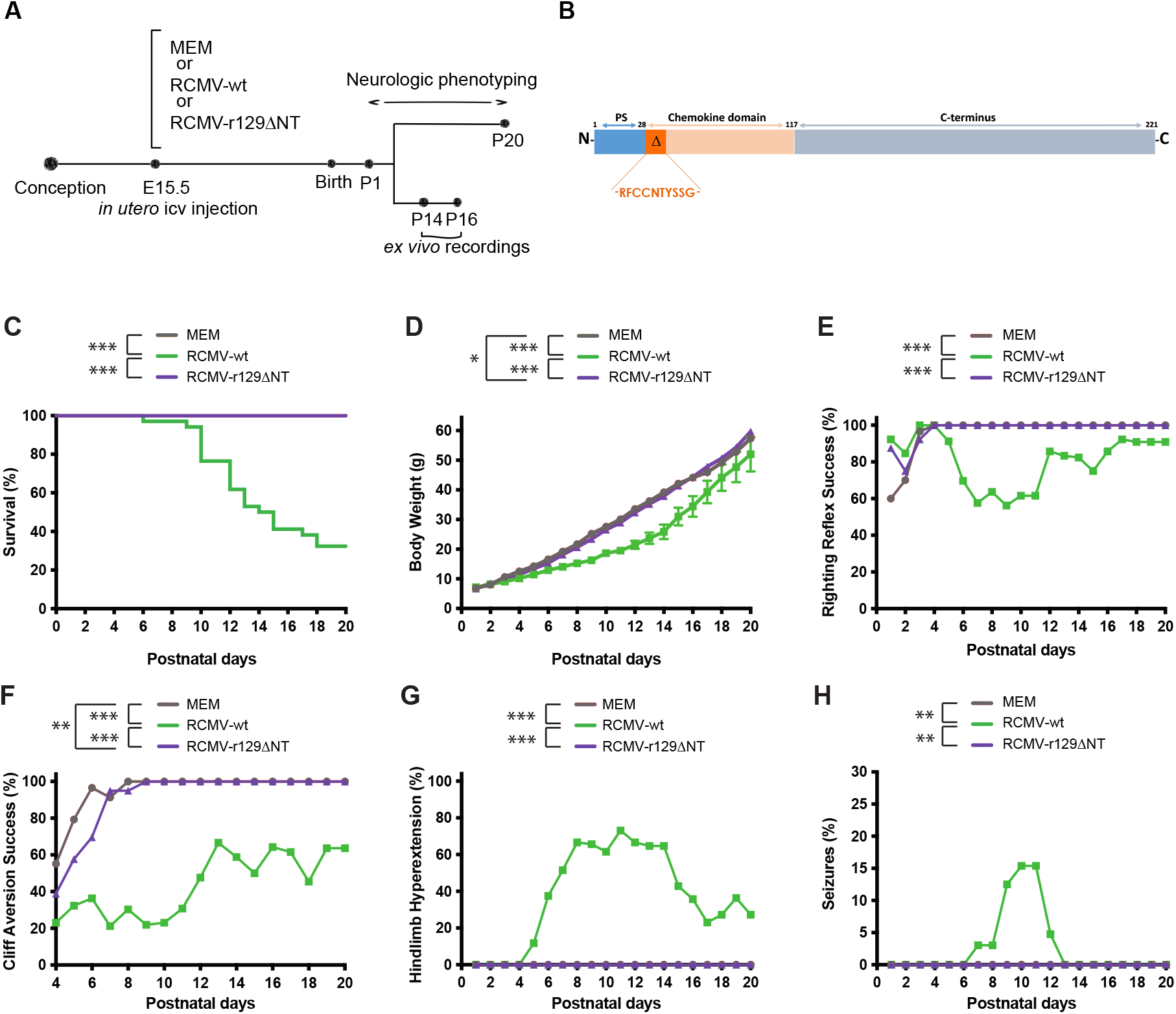
Infection of the fetal brain with rat CMV encoding a mutant RCK3 protein with a deleted CC chemokine motif (RCK3-ΔNT) is not pathogenic in vivo. (A) Experimental procedure and timeline. (B) Schematic view of the RCK3 protein. PS: peptide signal. Δ: the amino acid sequon deleted in RCK3-ΔNT is indicated. (C-H) Daily phenotypic evaluations in the three first postnatal weeks (postnatal days P1-P20) of (C) survival, (D) body weight, (E) success to the righting reflex test, (F) success to the cliff aversion test, (G) detection of hindlimb hyperextension, and (H) tonic-clonic epileptic seizures, in pups subjected to intracerebroventricular (icv) injection at embryonic day E15.5 of either wild-type CMV (RCMV-wt; n = 34, 3 litters), or mutant CMV (RCMV-r129ΔNT; n = 26, 3 litters), or the vehicle only (MEM; n = 29, 3 litters). Sex ratio did not differ significantly between the three groups at birth (p = 0.3529, Fisher’s exact test, two-tailed).

### RCMV encoding mutant RCK3 with a deleted CC chemokine motif exerts inhibitory effects on the postnatal consequences of wild-type RCMV

RCK3-ΔNT had been shown to exert a dominant-negative effect on the chemotaxis properties of wt RCK3 *in vitro* (Vomaske et al., 2012). We thus decided to co-infect the fetal brains with the wt and mutant viruses at E15.5 simultaneously. Three experimental conditions were used, in which different amounts (pfu) of mutant RCMV-r129ΔNT were administered while keeping the dose of RCMV-wt unchanged (Figure 2A). A dramatic and near-perfect prevention against all postnatal phenotypes classically caused by the *in utero* infection with RCMV-wt was obtained when five times more infectious dose of mutant RCMV-r129ΔNT than RCMV-wt was used (Figure 2B-G). In line with the expected titration effect of a dominant-negative effect, weaker rescue was obtained with twice less infectious dose of RCMV-r129ΔNT than RCMV-wt, while intermediate rescue was seen when equal infectious doses of both viruses were co-injected (Figure 2B-G).

**Figure 2.**
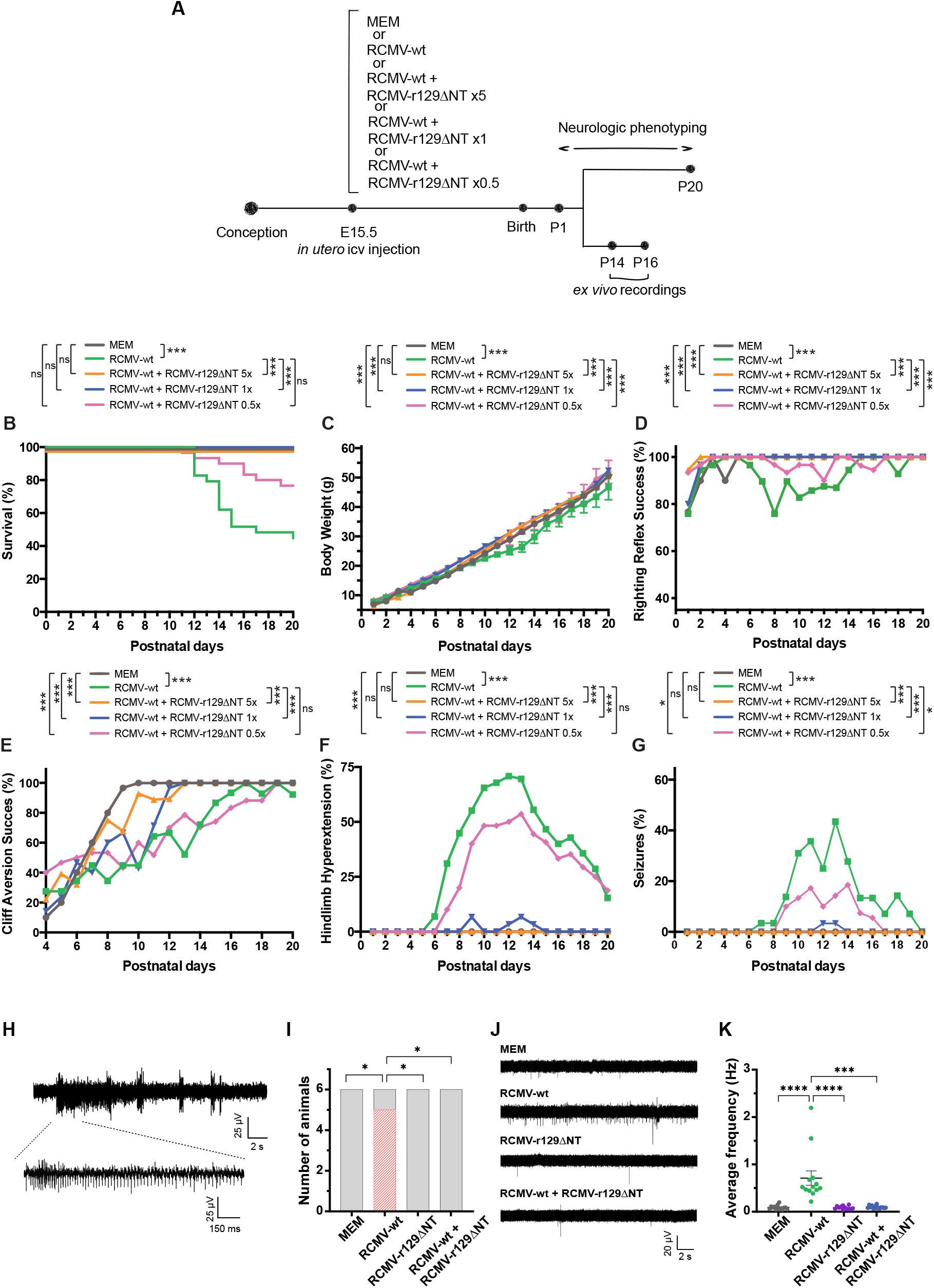
Infection of the fetal brain with mutant dominant-negative RCMV-r129ΔNT protects against the postnatal phenotypes and the epileptiform activity caused by RCMV-wt. (A) Experimental procedure and timeline. (B-G) Daily phenotypic evaluations in the three first postnatal weeks (postnatal days P1-P20) of (B) survival, (C) body weight, (D) success to the righting reflex test, (E) success to the cliff aversion test, (F) detection of hindlimb hyperextension, and (G) tonic-clonic epileptic seizures, in pups subjected to intracerebroventricular (icv) injection at embryonic day E15.5 of either wild-type CMV (RCMV-wt; n = 29, 3 litters), or wild-type (wt) CMV with various doses (x, compared to the dose of wt virus) of mutant CMV (RCMV-wt + RCMV-r129ΔNT 5× n = 28, 3 litters; RCMV-wt + RCMV-r129ΔNT 1x; n = 30, 3 litters; RCMV-wt + RCMV-r129ΔNT 0.5x; n = 30, 3 litters), or the vehicle only (MEM; n = 30, 3 litters). Sex ratio did not differ significantly between the five groups at birth (p = 0.3941, Fisher’s exact test, two-tailed). (H-K) Extracellular electrophysiological recordings within neocortical layer IV at postnatal days P14-P16 in brain slices from rat pups subjected to intracerebroventricular (icv) injection at embryonic day E15.5 of either wild-type CMV (RCMV-wt; n = 6, 3 litters), or mutant CMV (RCMV-r129ΔNT; n = n = 6, 2 litters), or equal doses of wild-type and mutant RCMVs together (RCMV-wt + RCMV-r129ΔNT; n = n = 6, 2 litters), or the vehicle only (MEM; n = 6, 2 litters). Sex ratio did not differ between the four groups at birth (p > 0.9999, Fisher’s exact test, two-tailed). (H) Representative local field potential (LFP) trace of a spontaneous ictal-like event recorded at P15 in brain slice from a rat pup infected with RCMV-wt *in utero*. The ictal-like event is also shown at an expanded timescale. (I) Number of pups exhibiting ictal-like events (dashed red); note that such events were detected only in the RCMV-wt condition. *: p < 0.05. Fisher’s exact test, two-tailed, with pairwise comparisons and Benjamini-Hochberg correction. (J) Representative LFP traces of spontaneous interictal-like events recorded in brain slices in the four conditions. (K) Summary data for averaged spike frequencies for interictal-like events in control condition (MEM, n = 14 slices from 6 pups (2 litters)), after infection with either of wild-type CMV (RCMV-wt, n = 13 slices from 6 pups (3 litters)) or mutant CMV (RCMV-r129ΔNT, n = 12 slices from 6 pups (2 litters)), and after co-infection with both wild-type and mutant CMVs (RCMV-wt + RCMV-r129ΔNT, n = 14 slices from 6 pups (2 litters)). ^****^: p < 0.0001 ^***^: p < 0.001. Kruskal-Wallis test with Dunn’s multiple comparisons test.

We then tested whether co-infection of RCMV-r129ΔNT with RCMV-wt also exerted its long-lasting rescuing effects at the level of neuronal networks. In line with our recently reported data in this model (Tarhini et al., 2024), extracellular electrophysiological recordings performed in layer IV of somatosensory cortices taken from pups aged 14-16 days (P14-P16) revealed spontaneous ictal-like events in brain slices from five out of the six RCMV-wt-infected pups, compared with the lack of any such epileptiform activity in six control (MEM-injected) pups (Figure 2H,I) (Figure S1). In sharp contrast with RCMV-wt, and in line with the phenotyping data *in vivo*, none of the slices from six RCMV-r129ΔNT-infected pups show any ictal-like activity upon recordings (p = 0.0304) (Figure 2I). Also, the significantly increased (p < 10^-4^) mean frequency of spontaneous interictal-like events detected in slices from RCMV-wt-infected pups (0.7105 Hz +/- 0.1514) compared to control slices (0.0895 Hz +/- 0.01158), was not observed in slices from RCMV-r129ΔNT-infected pups (0.0863 Hz +/- 0.001) (p < 10^-4^, compared to RCMV-wt-infected pups) (Figure 2J,K) (Figure S1). Importantly also, the epileptiform activity recorded in P14-P16 pups after RCMV-wt infection was strongly inhibited after co-infection of RCMV-r129ΔNT with RCMV-wt *in utero*: no ictal-like activity was ever detected in slices from six co-infected pups (p = 0.0304, compared to pups infected with RCMV-wt only) (Figure 2I), and the frequency of interictal-like events (0.0968 Hz +/- 0.0078) was significantly reduced (p = 0.0002) (Figure 2J,K) (Figure S1).

### The detrimental in vivo consequences of wild-type RCMV infection can also be counteracted with pre- or post-infection with the mutant RCMV

Taken together, the *in vivo* and *ex vivo* effects depicted above showed that simultaneous co-infection of the rat fetal brain with RCMV encoding mutant, dominant-negative RCK3 chemokine precluded the future emergence of the severe postnatal phenotypes caused by RCMV encoding wt RCK3, thus confirming the crucial role of RCK3 in CMV-related neuropathogenesis. This also indicated that the targeting of the virus-specific RCK3 chemokine might be beneficial in the context of congenital CMV infection, raising potential translational avenues. In this perspective, we decided to test whether the dominant-negative RCMV-r129ΔNT would also operate when administered not at the same time as RCMV-wt, but in a sequential way. To this aim, icv injection of mutant RCMV-r129ΔNT in the fetal brain was done at E14.5, hence 24h. before the injection, at E15.5, of an equal amount (pfu) of RCMV-wt (Figure 3A). As with the simultaneous co-injections described above, prior infection of the fetal brain with RCMV-r129ΔNT dramatically prevented against the severe postnatal consequences of RCMV-wt infection (Figure 3B-G). Remarkably, it was also possible to efficiently combat against the emergence of these severe postnatal phenotypes by administering RCMV-r129ΔNT icv not before or simultaneously as depicted before, but at E16.5, 24h after the injection of equal amount of RCMV-wt done at E15.5 (Figure 3B-G).

**Figure 3.**
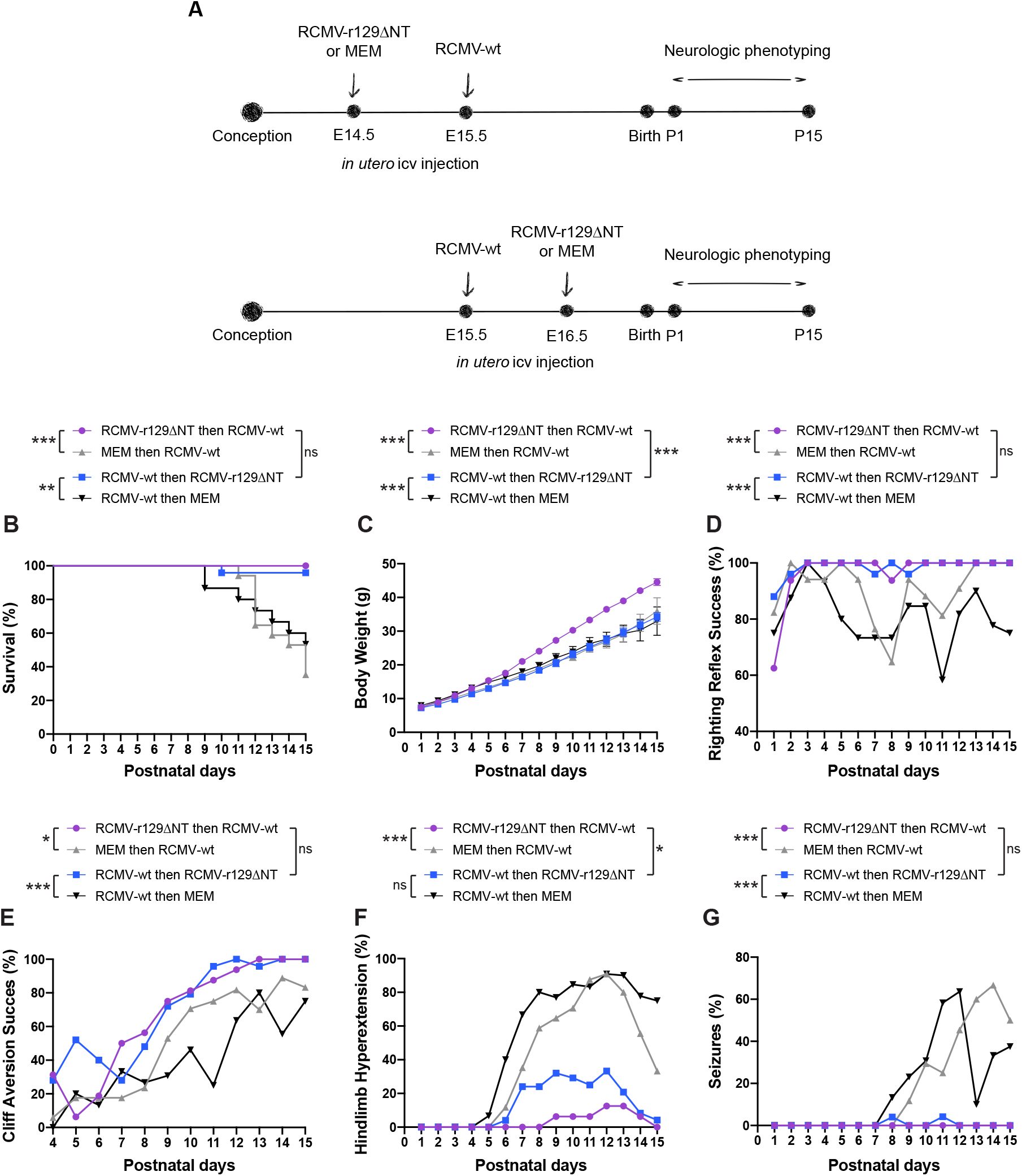
The deleterious postnatal phenotypes caused by RCMV-wt infection of the fetal brain can also be prevented by prior or by post infection with the mutant RCMV-r129ΔNT. (A) Experimental procedure and timeline. Top: E14.5/E15.5 consecutive icv injections. Bottom: E15.5/E16.5 consecutive icv injections. (B-G) Daily phenotypic evaluations in the two first postnatal weeks (postnatal days P1-P15) of (B) survival, (C) body weight, (D) success to the righting reflex test, (E) success to the cliff aversion test, (F) detection of hindlimb hyperextension, and (G) tonic-clonic epileptic seizures, in pups subjected to either of: intracerebroventricular (icv) injection at embryonic day E14.5 of mutant CMV followed by icv injection at E15.5 of wild-type CMV (RCMV-r129ΔNT then RCMV-wt; n = 16, 3 litters); icv injection at embryonic day E14.5 of MEM followed by icv injection at E15.5 of wild-type CMV (MEM then RCMV-wt; n = 17, 2 litters); icv injection at E15.5 of wild-type CMV followed by icv injection at embryonic day E16.5 of mutant CMV (RCMV-wt then RCMV-r129ΔNT; n = 25, 3 litters); icv injection at E15.5 of wild-type CMV followed by icv injection at embryonic day E16.5 of MEM (RCMV-wt then MEM; n = 15, 2 litters). Sex ratio did not differ significantly between the four groups at birth (p = 0.6059, Fisher’s exact test, two-tailed).

## Discussion

Here, we show that a chemokine encoded by rat CMV, RCK3, is critically involved in the neurodevelopmental pathogenesis caused by infection of the rat fetal brain. A recombinant rat CMV encoding a mutant isoform of RCK3 bearing a deletion in the CC chemokine domain, failed to induce the deleterious postnatal phenotypes caused by wt rat CMV *in vivo* and *ex vivo*. Furthermore, rat CMV with dominant-negative mutant RCK3 exerted an inhibitory effect on wt rat CMV *in vivo*; this effect was robust enough to allow near-perfect prevention against the postnatal phenotypes and was obtained whether the infection with mutant rat CMV occurred before, at the same time, or after infection with wt rat CMV.

### RCK3 chemokine and neurodevelopmental pathogenesis

The pathogenic mechanisms driven by RCK3-related chemotaxis activity are likely multiple and complicated. CMV encodes several proteins including RCK3 that might disturb key neurodevelopmental events by interfering with host chemokine/chemokine receptor systems (Pontejo & Murphy, 2017). RCK3 binds several host CC chemokine receptors (CCRs) (Vomaske et al., 2012). Hence, RCK3 could interfere with the control of neuronal autophagy, which can be driven by microglial chemokines CCL3-5 *via* the neuronal RCK3 receptor CCR5 (Festa et al., 2023). Interestingly, CCR2, another CC chemokine receptor, has been associated with neuropathological processes in the context of viral encephalitis (Käufer et al., 2018). Also, crucial neuronal oscillations such as giant depolarizing potentials that shape neuronal circuits during development, or the proper laminar destination of Lhx6^+^ interneurons in the neocortex, are influenced by various chemokines/chemokine receptors (Bertot et al., 2019; Tiveron et al., 2006). As a matter of fact, our electrophysiological recordings of neocortical networks revealed abnormal, epileptiform activity after prior infection of the fetal brain with wt CMV, but not with mutant rat CMV coding for the mutant RCK3. Not exclusively, RCK3 might be involved in rat CMV neuropathogenesis by impacting on the recruitment and on the distribution of immune cells in the infected developing brains. The alteration of fetal microglia (Cloarec et al., 2018; Tarhini et al., 2024) and various non-resident immune cells such as memory CD8^+^ T-cells (Brizić et al., 2018) and natural killer (NK) and type 1 innate lymphoid cells (Kveštak et al., 2021) have already been implicated in rodent models of congenital CMV infection.

### Dominant-negative approaches for CMV and other viral infections

In the present study, the coding sequence of dominant-negative isoform of RCK3 inserted in the rat CMV genome acted upon infection of the corresponding mutant virus in host cells, revealing a key pathogenic role for RCK3. Mutations with dominant-negative effects encode proteins that nullify the function of the wild-type isoform (Herskowitz, 1987). Naturally-occurring dominant-negative mutations have been identified in a plethora of genetic disorders, and methods based on dominant-negative effects have long been proposed as a way to target proteins of interest involved in key physiological or pathological processes. Dominant-negative proteins have been widely used to decipher the function of herpesvirus genes. The use of dominant-negative effect against viral infection was initially reported with a truncated isoform of herpes simplex virus-1 (HSV-1) transcription activator VP16 providing resistance against infection (Friedman et al., 1988): this led to the principle of intracellular immunization for generating antiviral effects (Baltimore, 1988), which has then been employed not only in the context of herpes viruses (Augustinova et al., 2004; Mühlbach et al., 2009) and of CMV notably (Čičin-Šain et al., 2008; Popa et al., 2010; Rupp et al., 2007), but also of human immunodeficiency virus (HIV-1) (Inubushi et al., 1998; Trono et al., 1989). Deciphering how RCK3-ΔNT exerted its dominant-negative effect on RCK3-wt in the simultaneous and consecutive co-infection rescue assays *in vivo* is beyond the scope of the present study. As classically seen for secreted proteins with dominant-negative isoforms (Haslund et al., 2018; Riva et al., 2024), mutant RCK3-ΔNT might act intracellularly in co-infected cells by impairing the secretion of RCK3-wt and possibly also by favoring its degradation, likely through the formation of wt/mutant dimers. Extracellular interference, for instance *via* competition for RCK3 receptors binding sites, might be involved as well.

Methods based on genetic interference represent starting points for the design of novel antiviral strategies (Tanner et al., 2016), as the targeting of viral proteins would limit the risk to off-targets effects. Very recently, interference based on the co-infection of an engineered virus with the corresponding wt virus was applied to HSV-1 in mice to spread a foreign gene to the wt viruses in the infected cells *via* CRISPR-mediated homologous recombination (Walter et al., 2024). While our study was based on a simpler dominant-negative approach, the design and use of engineered herpesviruses to combat against themselves was expanded here to a congenital context: dramatic prevention of the postnatal consequences of CMV infection of the fetal brain was obtained by prenatal co-infection assays with a mutant CMV encoding a dominant-negative isoform of viral chemokine RCK3. This did not only demonstrate a crucial pathogenic role for RCK3, but also indicated that approaches aimed at selectively targeting viral chemokines *in utero* could be very efficient against the detrimental consequences of congenital CMV infection, and provided a proof-of-concept for the possible use *in vivo* of dominant-negative interference approaches targeting the CC chemokine domain of UL128, the functional homolog of RCK3 encoded by human CMV. It was already suggested that mutation in the CC chemokine domain of GP129, the homolog of UL128 in guinea pig CMV, had a role in viral pathogenicity irrespective of pentameric complex formation (Coleman et al., 2017). The CC chemokine domain of UL128, the functional homolog of RCK3, would thus represent a promising target for the engineering of human recombinant CMVs with dominant-negative effects.

## Author contributions

SB performed most experiments and data analyzes, co-decided on the overall strategy and co-directed the follow-up of the experiments with PS. ST performed all electrophysiological experiments and participated in data analyzes and in manuscript writing. EB supervised and performed all *in utero* intracerebroventricular infections. SS designed and performed phenotypic statistical analyzes with ST. TS performed production and purification of the rat CMV strains. DNS provided and engineered the recombinant rat CMV strains. NB supervised electrophysiological recordings and analyses. HL participated in study design and strategy. PS co-decided on the overall strategy and co-directed the follow-up of experiments with SB, and wrote the manuscript with SB and with participation of ST. All authors contributed the final version of the manuscript.

## Acknowledgments

We thank Séverine Corby and all members of the INMED (Mediterranean Institute of Neurobiology) animal core facilities.

## Supplementary Material

Figure S1

