## Supplementary material for "Virus-against-virus dominant-negative interference strategy targeting a viral CC chemokine prevents cytomegalovirus-related neurodevelopmental pathogenesis": Figure S1

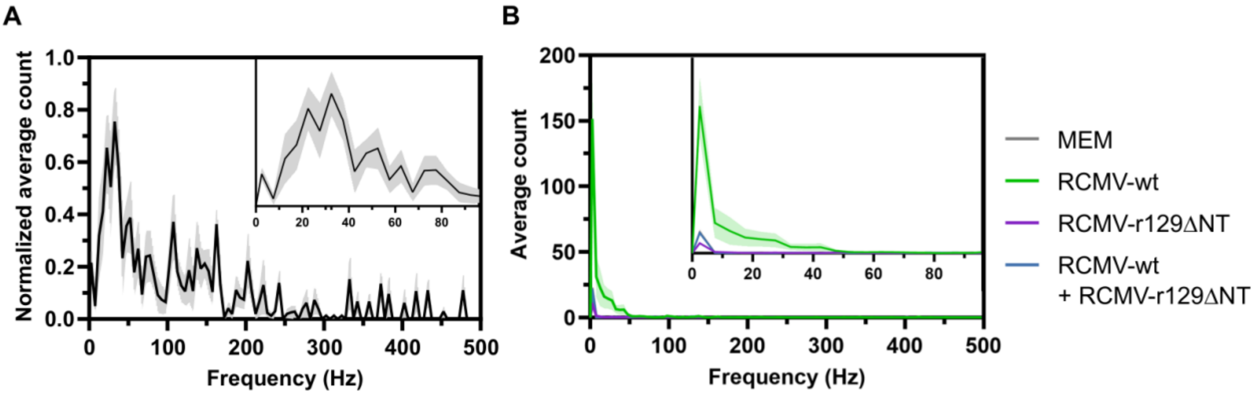


Figure S1. *Distribution frequency of ictal-like (A) and interictal-like (B) events*.
Extracellular electrophysiological recordings within neocortical layer IV at postnatal days P14-P16 in brain slices from rat pups subjected to intracerebroventricular (icv) injection at embryonic day E15.5 of either wild-type CMV (RCMV-wt; n = 6, 3 litters), or mutant CMV (RCMV-r129ΔNT; n = 6, 2 litters), or equal doses of wild-type and mutant RCMVs together (RCMV-wt + RCMV-r129ΔNT; n = 6, 2 litters), or the vehicle only (MEM; n = 6, 2 litters). Sex ratio did not differ between the four groups at birth ($p > 0.9999$, Fisher's exact test, two-tailed).

(A) Mean frequency distribution of spikes for ictal-like events (n = 6 events from 5 out of 6 RCMV-wt-infected pups). Bin size was set at 5 Hz. Normalized average count values corresponding to the middles of each bin are connected with a solid line. A zoom-in of the initial part (0-100 Hz) of the frequency distribution is shown on the right corner.

(B) Mean frequency distribution of spikes for interictal-like events. RCMV-wt-infected pups (RCMV-wt): green trace, n = 13 slices from six pups; RCMV-r129ΔNT-infected pups (RCMV-r129ΔNT): purple trace, n = 12 slices from six pups; RCMV-wt + RCMV-r129ΔNT co-infected pups (RCMV-wt + RCMV-r129ΔNT): blue trace, n = 14 slices from six pups; non-infected, MEM-injected pups (MEM): grey trace, n = 14 slices from six pups. A zoom-in of the initial part (0-100 Hz) of the frequency distribution is shown on the right corner.
